# Single-cell joint detection of chromatin occupancy and transcriptome enables higher-dimensional epigenomic reconstructions

**DOI:** 10.1101/2020.10.15.339226

**Authors:** Haiqing Xiong, Yingjie Luo, Qianhao Wang, Xianhong Yu, Aibin He

## Abstract

Deciphering mechanisms in cell fate decisions requires single-cell holistic reconstructions of multi-dimensional epigenome in transcriptional regulation. Here we develop CoTECH, a combinatorial barcoding method allowing for high-throughput single-cell joint detection of chromatin occupancy and transcriptome. First, we used CoTECH to examine bivalent histone marks (H3K4me3 and H3K27me3) with transcription from naïve to primed mouse embryonic stem cells. Concurrent bivalent marks in pseudo-single cells linked via transcriptome were computationally derived, resolving pseudotemporal bivalency trajectories and disentangling a context-specific interplay between H3K4me3/H3K27me3 and transcription level. Next, CoTECH with H3K27ac, an active enhancer marker, revealed the regulatory basis of endothelial-to-hematopoietic transition in two waves of hematopoietic cells and distinctive enhancer-gene linking schemes guiding hemogenic endothelial cell (HEC) emergence, indicating a unique epigenetic control of transcriptional regulation for hematopoietic stem cell priming. Together, CoTECH provides an efficient framework for single-cell co-assay of chromatin occupancy and transcription, thus, enabling higher-dimensional epigenomic reconstructions.

## INTRODUCTION

Single-cell transcriptome sequencing (scRNA-seq) has revolutionized our understanding of cellular heterogeneity in physiological and pathological biological processes (Bowling et al., 2020; Cao et al., 2019; Klein et al., 2015; Macosko et al., 2015; Patel et al., 2014; Picelli et al., 2013; Pijuan-Sala et al., 2019; Regev et al., 2017; Spanjaard et al., 2018; Tabula Muris et al., 2018; Tang et al., 2009; Zhou et al., 2016). The transcriptome and cell identities of different cell types/states with the same DNA are determined by multilayers of epigenetic information. Recently, technologies for single-cell profiling of multi-dimensional chromatin states have been developed, such as various scChIP-seq techniques for DNA-binding proteins and histone modifications (Ai et al., 2019; Kaya-Okur et al., 2019; Ku et al., 2019; Rotem et al., 2015; Wang et al., 2019), scATAC-seq for chromatin accessibility (Buenrostro et al., 2015), MNase-seq for nucleosome positioning (Lai et al., 2018), scBS-seq for DNA methylome (Guo et al., 2015) and scHiC for higher-order chromatin structure (Nagano et al., 2013). Although these methods measure multiple modalities of single cells, each provides only specific layers of cellular heterogeneities. To build connections across these layers in single cells, innovative computational platforms emerge to integrate unpaired single-cell omics datasets and project different molecular information into a common latent space (Efremova and Teichmann, 2020; Stuart et al., 2019). However, existing strategies require priori knowledge-based correspondence across multimodal omics datasets from different experiments, limiting the ability to reconstruct an accurate view of functional relationship between different modalities of epigenomic features and gene expression as well as the crosstalk between different epigenetic layers.

More recently, unlike those profiling molecular layers one at a time, new approaches have been developed to experimentally link the transcriptome and epigenome by simultaneously measuring multi-omics in the same single cells, making it possible to precisely analyze single-cell-resolved epigenomic regulation of gene expression and cell fate decisions. For example, several various single-cell co-assays (combined scATAC-and scRNA-seq) for joint analysis of accessible chromatin and gene expression have been developed, permitting inference of the correlation between cis-regulatory elements and putative target genes (Cao et al., 2018; Chen et al., 2019; Clark et al., 2018; Zhu et al., 2019).

Apart from chromatin accessibility, other important molecular layers of the epigenome are covalent modifications to histones and chromatin occupancy of DNA binding proteins, providing a critical guidance for determining transcriptional outcomes (Park, 2009). Two elegant studies demonstrated the proof-of-concept application of simultaneously quantifying of protein–DNA interactions and transcriptome in single cells by scDam&T-seq and scCC (Moudgil et al., 2020; Rooijers et al., 2019). Both methods rely on a transgene expressing transcription factors or chromatin binding proteins tethered to *Escherichia coli* DNA adenine methyltransferase (Dam) or piggyBac transposase for scDam&T-seq and scCC, respectively, limiting the likelihood of wide adoption and the potential implementations, in particular for studying clinical human samples. Moreover, the same strategy cannot measure various histone modifications in single cells.

Here, we develop a high-throughput method for single-cell joint detection of chromatin occupancy and transcriptome. This approach, named CoTECH (<u>co</u>mbined assay of <u>t</u>ranscriptome and <u>e</u>nriched <u>c</u>hromatin binding), adopts a combinatorial indexing strategy (Cusanovich et al., 2015; Wang et al., 2019) to enrich chromatin fragments of interest as reported in CoBATCH in combination with a modified Smart-seq2 procedure (Picelli et al., 2014) to simultaneously capture the 3, mRNA profiles in the same single cells. To demonstrate the utility of CoTECH, we used it to study the relationship between multiple histone modifications and gene expression in mouse embryonic stem cells (mESCs) and the regulatory basis of endothelial-to-hematopoietic transition (EHT) in two waves of hematopoietic cells. Experimentally linking chromatin occupancy to transcriptional outputs and inferred molecular association between multimodal omics datasets made possible by CoTECH enables reconstructions of higher dimensional epigenomic landscape, providing new insights into epigenome-centric gene regulation and cellular heterogeneity in many biological processes.

## RESULTS

### Designing CoTECH for joint profiling of transcriptome and chromatin occupancy in single cells

To accurately align the multi-layered omics information within the same cell, we developed CoTECH, a method for a combinatorial indexing-based single-cell co-assay for transcriptome and chromatin occupancy. Profiling chromatin parts in CoTECH is based on our previously well-established CoBATCH (Wang et al., 2019) with a few modifications that allow simultaneously measuring mRNA by the following steps (Figures 1 and S1; Table S1; STAR Methods): (i) Single cells on concanavalin A coated beads are permeabilized and incubated with primary and secondary antibodies. (ii) Cells bound with the antibody are distributed into a 96-well plate with protein A-fused Tn5 transposon (PAT) assembled with a unique combination of T5 and T7 barcodes in each well. (iii) *in situ* tagmentation of chromatin bound by the protein-antibody complex. (iv) Reverse transcription of mRNA using oligo dT primer bearing an 8-bp well-specific first-round RNA barcode and an 8-bp unique molecular identifier (UMI), and template switch using TSO oligo. (v) Pooling and re-distribution of cells into multiple 96-/384-well plates. (vi) Pre-amplification of cDNA and tagmented chromatin segments. (vii) DNA purification and split into two halves for RNA- and DNA-devoted library preparation, separately. (viii) For RNA-part, pre-amplified full-length cDNA is tagmented by transposon Tn5, and 3, end of the tagmented cDNA is amplified by introducing a second round of RNA index. (ix) For DNA-part, chromatin DNA is amplified by using a pair of common adaptor primers in the first round and a pair of well-specific indexing primers in the second round of PCR.

**Figure 1.**
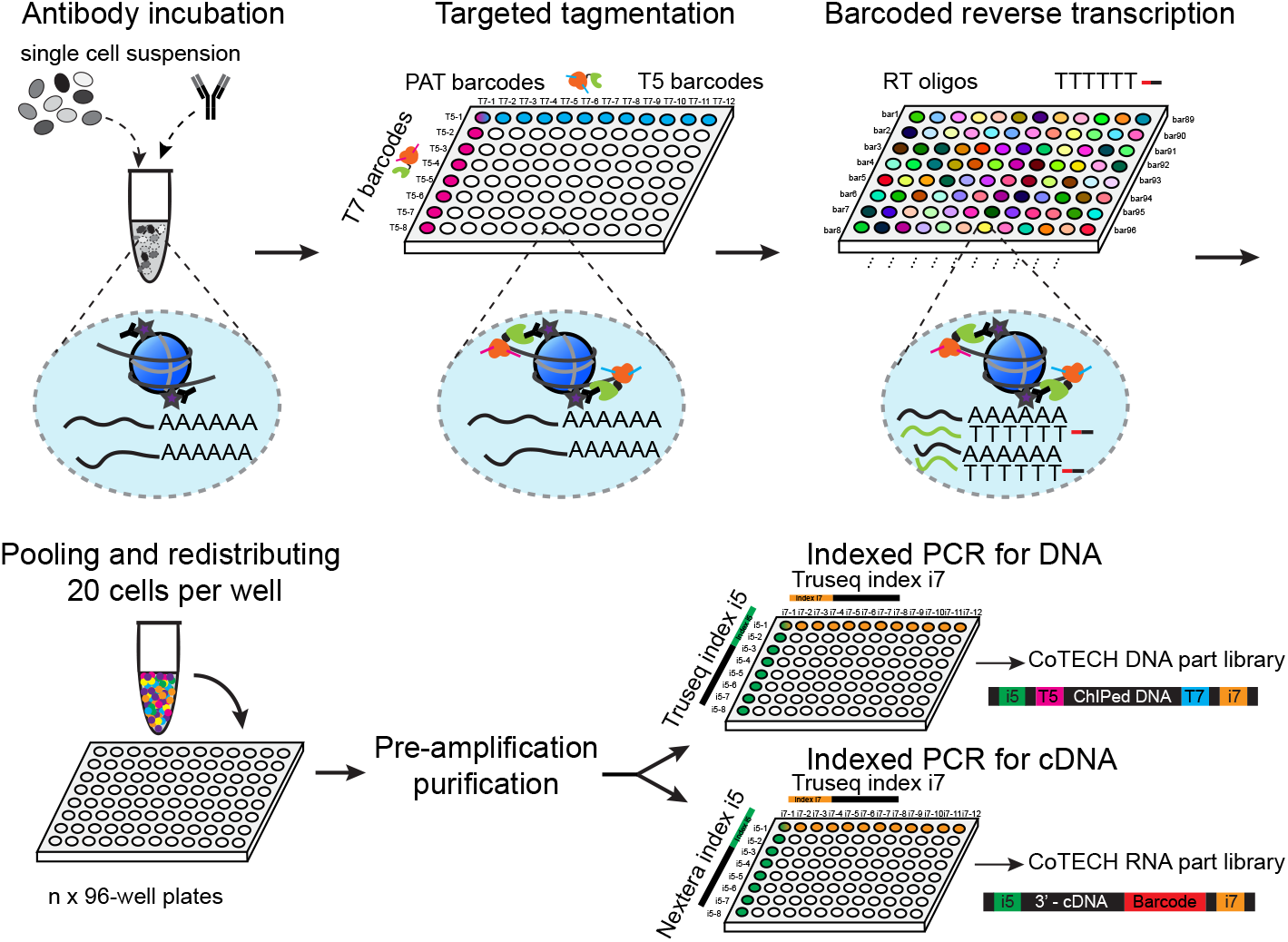
Schematic design of CoTECH. Single-cell suspension was incubated with antibody before FACS sorting 2,000 cells into each well with a unique combination of barcoded adaptors with Tn5 fused to protein A as well as barcoded primers for reverse transcription (RT). After tagmentation and RT, cells were pooled together and redistributed into a number of secondary 96-well plates for pre-amplification of both cDNA and genomic DNA parts. Samples were purified with AMPure XP beads and divided into two halves for preparation of DNA and RNA libraries, respectively.

To assess the ability of CoTECH based on combinatorial indexing to distinguish single cells, we designed a proof-of-principle human-mouse “species-mixing” experiment on H3K27ac and RNA in 1:1 mixed HEK293T and NIH 3T3 cells. We found that the collision rate (the proportion of cells coincidentally receiving the same barcode combination) was no more than 4% for both DNA and RNA portions, confirming a successful indexing strategy for distinguishing single cells (Figures 2A and S2A-S2C).

**Figure 2.**
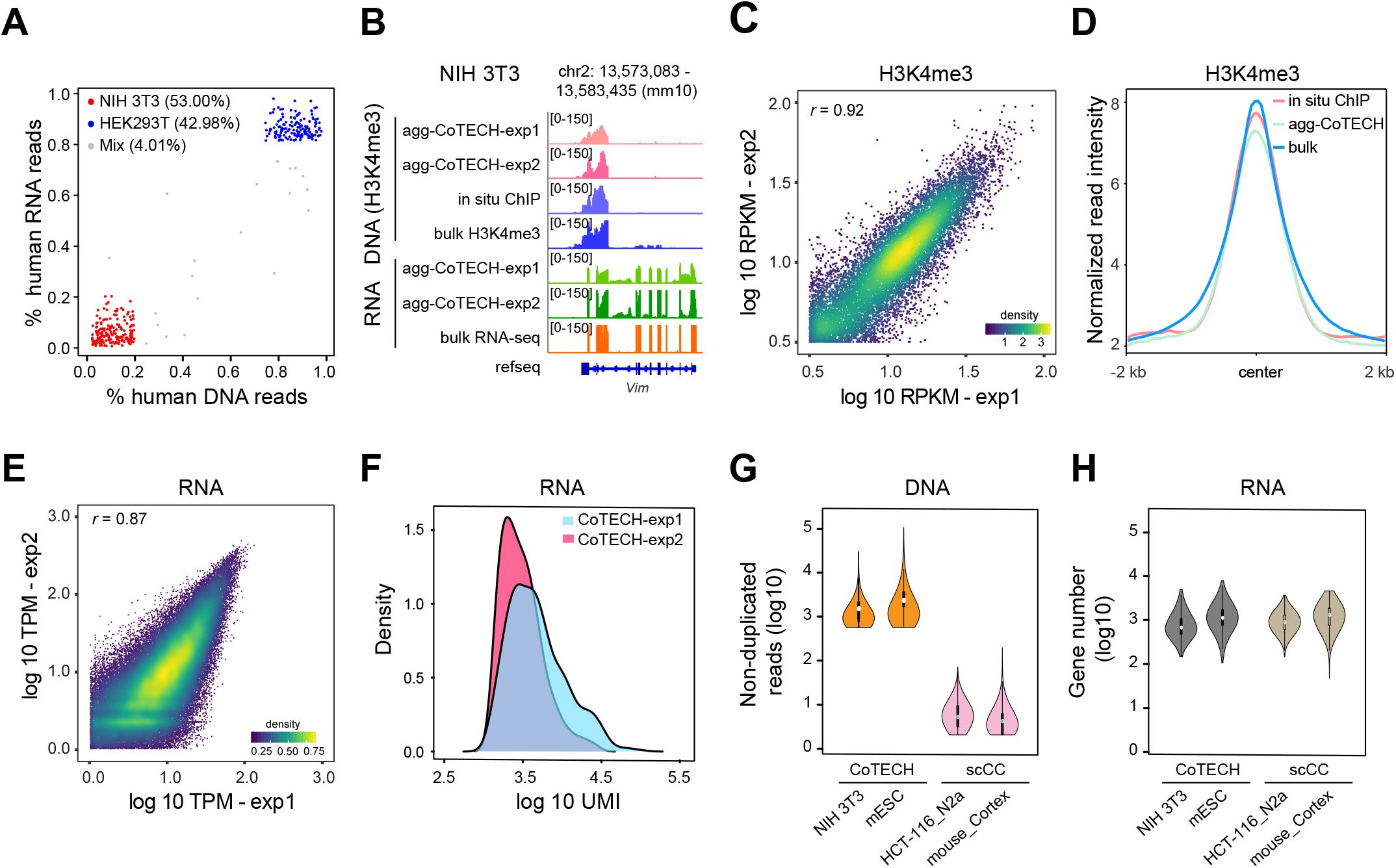
Assessing the ability of CoTECH to capture histone mark and transcription in single cells. (A) Scatter plot in the Human-mouse mix test for cells with both DNA and RNA profiles obtained. (B) Track view displaying both H3K4me3 and RNA signals on a representative locus in NIH 3T3. Bulk H3K4me3 ChIP-seq and RNA-seq data were downloaded from GSM1544520 and GSM970853, respectively. (C) Pearson correlation of H3K4me3 signals between two CoTECH experiments. Aggregate H3K4me3 signals were log10 normalized. (D) Average H3K4me3 signals from different datasets were plotted at the 2-kb flanking regions around the center of 25,311 peaks. Bulk data were from GSM1544520. (E) Track view showing Pearson correlation of transcriptome between two CoTECH experiments. The transcription level was normalized as TPM. (F) The distribution of detected UMI between two CoTECH experiments in NIH 3T3 cells (n = 844). (G-H) Violin plot showing the distribution of non-duplicated reads (DNA) (G), gene number (RNA) (H) in different datasets across CoTECH and scCC (Moudgil et al., 2020). Processed data were downloaded from GSE148448.

We next performed CoTECH experiments on H3K4me3 and RNA in NIH 3T3 cells to benchmark CoTECH and further evaluate the data quality. Reads from the RNA library were highly enriched in exonic regions as demonstrated in the regions of *Vim* gene loci from both aggregate CoTECH single-cell and bulk data (Figures 2B and S2D). Expectedly, we observed strong H3K4me3 signals at the promoter region of the representative gene for the corresponding DNA portion of CoTECH, low-input *in situ* ChIP-seq, and bulk ChIP-seq datasets (Figure 2B). For DNA portion of CoTECH experiments in NIH 3T3 cells, we acquired a median of 1,153 non-duplicated reads in 844 co-assayed cells, with a median FRiP (fraction of reads in peaks) of 62.9% (Figure S2E). Global analysis revealed a high correlation (Pearson correlation coefficient, 0.92) for the aggregate H3K4me3 signals between two CoTECH experiments. CoTECH DNA profiles markedly recapitulated those from in situ ChIP and bulk ChIP-seq at 25,311 peak regions (Figures 2C and 2D). CoTECH RNA profiles were also highly reproducible between two experiments (Pearson correlation coefficient, 0.87) (Figure 2E). The CoTECH experiments on NIH 3T3 yielded a median of 3,490 UMIs (unique molecular identifiers) per cell from 844 single cells (Figure 2F). We found 75% RNA reads mapped to the mouse genome, with 40% of those mapped to exons (Figure S2D). Compared with a recent high-throughput method in a different design, named scCC (Moudgil et al., 2020), using a transgene expressing an exogenous TF-transposase fusion for profiling RNA and TF-DNA interactions, CoTECH captured more non-duplicated reads per cell in DNA parts (~1,500 reads in COTECH versus ~ 10 reads in scCC) (Figures 2G, 2H and S2F). We also benchmarked our CoTECH data alongside other high-throughput multi-omics approaches (sci-CAR, Paired-seq and SNARE-seq)(Cao et al., 2018; Chen et al., 2019; Zhu et al., 2019) but for joint assay of RNA and accessible chromatin. We found that there were inappreciable differences in both the non-duplicated reads and gene numbers detected per cell, while CoTECH exhibited markedly highest UMIs than all of those (Figure S2G). Notably, optimizing the pre-amplification process for both DNA- and RNA-devoted parts in K562 cells, we significantly improved the detection sensitivity of DNA non-duplicated reads, yielding the median 3,023 reads (Figure S3). The optimized protocol was then applied to the following CoTECH experiments. Taken together, these results support that CoTECH is able to sufficiently capture both gene expression and histone modifications in the same cell.

### Profiling of bivalent histone marks in mESCs with CoTECH for reconstruction of the gene-regulatory relationship

Next, we sought to explore the relationship between histone modifications H3K4me3/H3K27me3 and transcription in single cells. After filtering out low-quality cells, a total of 6,993 single cells with both chromatin occupancy and transcriptome were obtained (3,907 cells for H3K4me3-RNA CoTECH and 3,086 cells for H3K27me3-RNA CoTECH), with a median of 1,506 non-duplicated reads and 7,568 UMIs per cell, for following analyses. Data quality in dual omics profiling was exemplified by specific signals from CoTECH and public bulk datasets on genomic tracks at the *Pou5f1* and *Gata2* gene loci, confirming the expected pattern for H3K4me3 (an active mark), H3K27me3 (a repressive mark) and transcription in mESCs (Figure 3A). CoTECH experiments in mESCs presented a high robustness and reproducibility without batch effect (Figures S4A-S4D). Using RNA profiles, cells were clustered into two subpopulations (C1 and C2) annotated as “ naïve mESCs” and “primed mESCs” according to the expression of pluripotency marker genes (Figures 3B, 3C and S4E). Independently, we also identified two clusters using either CoTECH H3K4me3 or H3K27me3. 68.7% and 82.6% of co-assayed single cells from RNA-H3K4me3 and RNA-H3K27me3 experiments exhibited concordant cluster assignment, respectively. Most discordant cells at the border of C1 and C2 likely reflected an asynchronous change between transcriptome and epigenome or suggested a need for considering multimodal omics information together for cell clustering or definition during cell fate transition (Figures 3D–3G).

**Figure 3.**
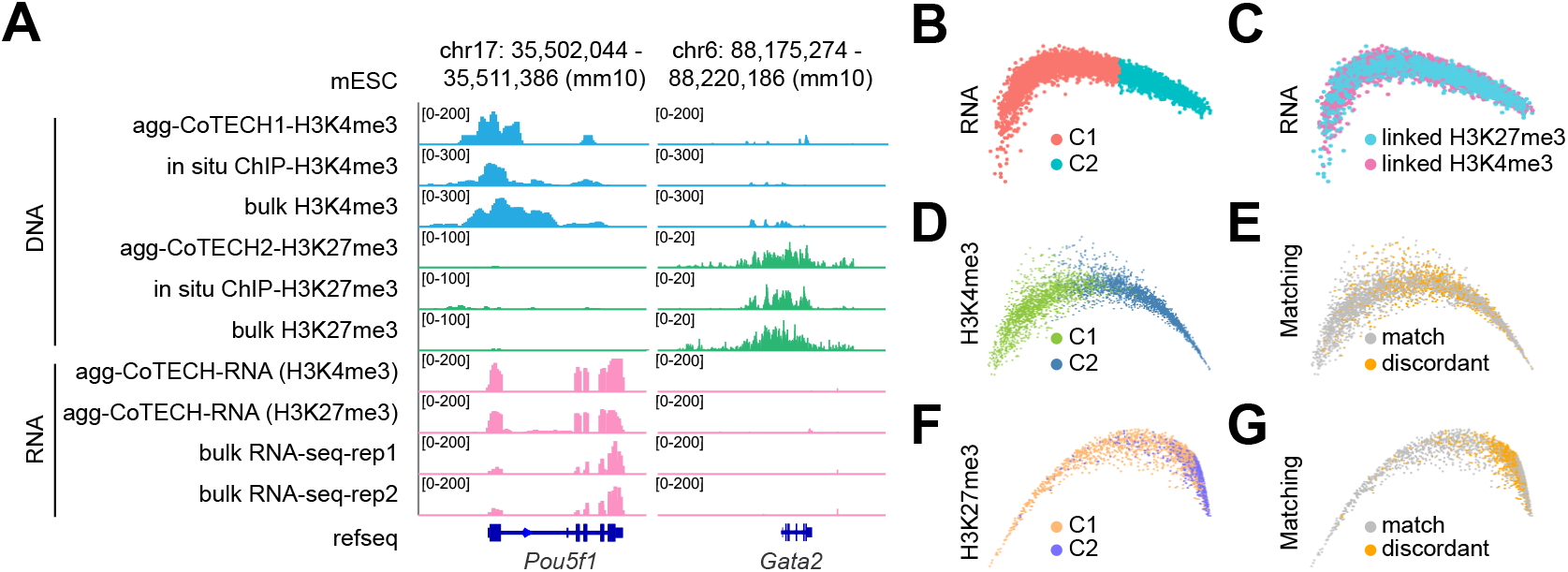
Single-cell joint detection of H3K4me3/H3K27me3 and transcription in mouse embryonic stem cells. (A) Track view displaying both H3K4me3, H3K27me3 signals and transcription (RNA) on the representative locus in mouse embryonic stem cells (mESCs). Bulk H3K4me3, H3K27me3 ChIP-seq and bulk RNA-seq data were downloaded from GSM1000124, GSM1000089, and GSM723776, respectively. (B-C) Dimension reduction of 6,993 single cells (both DNA and RNA detected) clustered by RNA profiles (B), and corresponding H3K4me3 or H3K27me3 profiles (C) in mESCs. (D-E) Diffusion map of 3,907 single cells (both DNA and RNA detected) clustered by H3K4me3 profiles (D) in mESCs. Cells in the same clusters in both H3K4me3 and RNA data are labelled with “match”, otherwise labelled with “discordant” in (E). (F-G) Diffusion map of 3,086 single cells (both DNA and RNA detected) clustered by H3K27me3 profiles (F) in mESCs. Cells in the same clusters in both H3K27me3 and RNA data are labelled with “match”, otherwise labelled with “discordant” in (G).

A predominant view has asserted that H3K4me3 as an active mark on chromatin plays an instructive role in transcriptional activation (Benayoun et al., 2014; Howe et al., 2017; Ruthenburg et al., 2007). Next, we capitalized on joint multi-omics information by CoTECH to revisit this question at single-cell and genome-wide spatial resolution. Indeed, in agreement with previous reports on bulk ChIP-seq data, aggregate CoTECH data in mESCs presented a positive correlation between the transcriptional level and H3K4me3 signals on both the regions of transcription start site (TSS) ± 5 kb (Pearson correlation coefficient; 0.75) and gene body ± 100 kb (Pearson correlation coefficient; 0.79) (Figure 4A) (Barski et al., 2007; Santos-Rosa et al., 2002; Wang et al., 2008). However, when examined at single cell resolution, this correlation was dramatically reduced to 0.05 at TSS regions (Figure 4B), despite several pluripotency genes such as *Pou5f1* and *Klf4* still displayed an overall positive consistency (Figure 4C). Although it is possible that the less significant correlation of H3K4me3 and gene expression at single cells might be related with the data sparsity, this result also potentially reflected the cell-to-cell asynchrony and variation on the molecular layers of transcriptome and epigenome.

**Figure 4.**
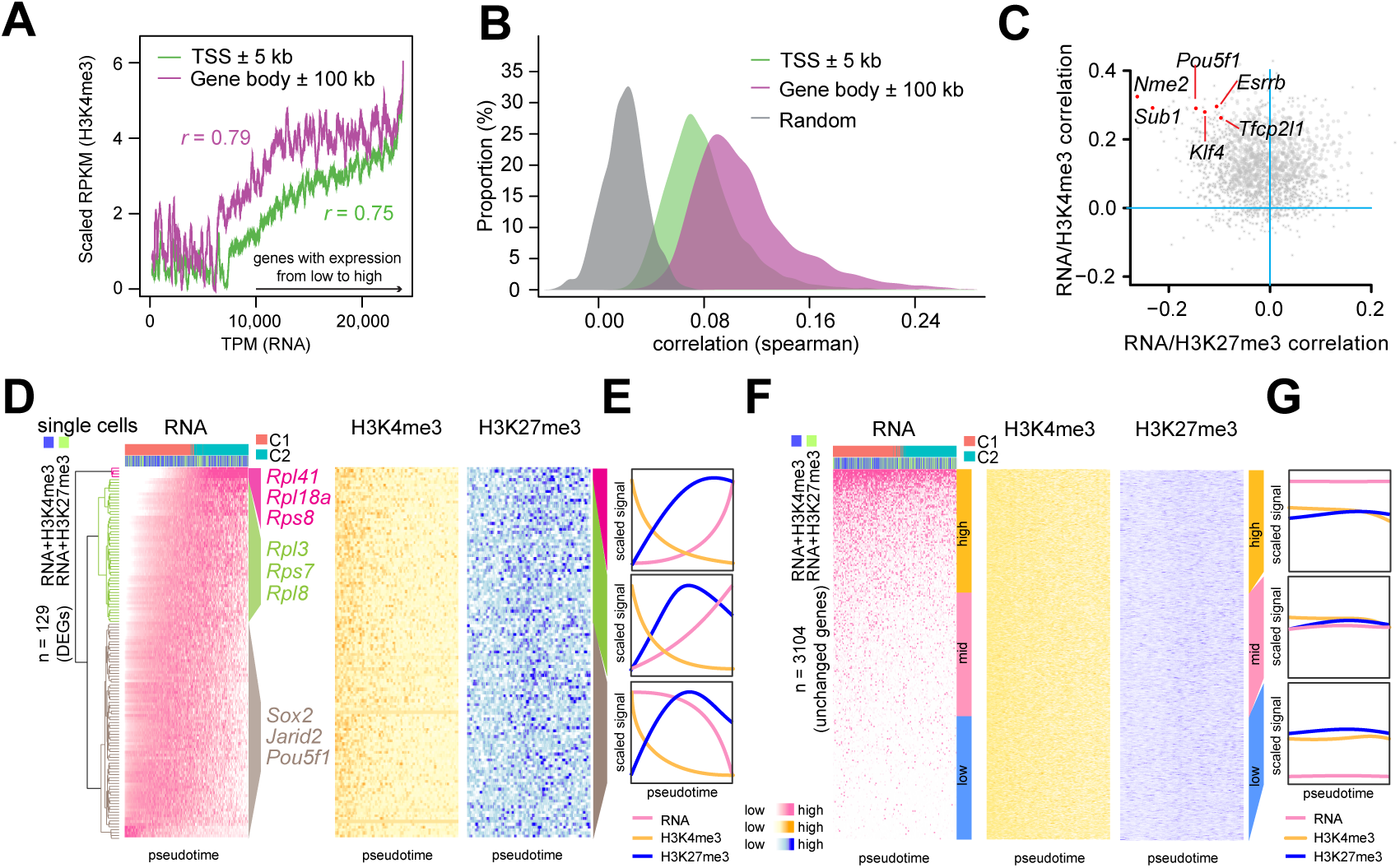
CoTECH reveals both coupling and uncoupling of dynamic histone modifications and transcription from naive to primed mESCs. (A) Moving average plots showing correlation between RNA and H3K4me3 signals at TSS ± 5 kb (green) and gene body ± 100 kb (magenta) using aggregate CoTECH data. (B) Distribution of correlation between RNA and H3K4me3 signals in single cells. (C) Scatter plots showing the spearman correlation coefficients of H3K27me3 versus RNA signals (x axis) and H3K4me3 versus RNA (y axis). Each dot represents one gene (top 2,000 variable genes were used). (D) Heatmap showing the normalized RNA signals of differentially expressed genes across 6,993 mESCs (left), normalized H3K4me3 (middle) signals in 3,907 cells and H3K27me3 (right) signals in 3,086 cells at gene body ± 100-kb regions. 129 differentially expressed genes were clustered to three clades. The columns represent single cells ordered by pseudotime. (E) Aggregate curves showing average RNA, H3K4me3, H3K27me3 signals in (D) of three gene classes as in (D). (F) Heatmap showing the normalized RNA signlas of differentially expressed genes across 6,993 mESCs (left), normalized H3K4me3 (middle) signals in 3,907 cells and H3K27me3 (right) signals in 3,086 cells at gene body ± 100 kb regions. 3,104 unchanged genes were divided into three groups based on the expression level. The columns represent single cells ordered by pseudotime. (G) Aggregate curves showing average RNA, H3K4me3, H3K27me3 signals in (F) of three gene classes as in (F).

We examined the dynamic change in transcription and H3K4me3/H3K27me3 from the naïve to primed mESCs. We ordered single cells based on RNA profiles along a pseudotime trajectory by Monocle 2 (Figures S4F and S4G). Out of most differentially expressed 129 genes, the pluripotency genes were downregulated and ribosomal-protein genes were upregulated along pseudotime, consistent with previously reported (Fortier et al., 2015). These genes in the single-cell matrix were clustered into 3 classes according to the dynamic expression pattern along pseudotime (Figures 4D and 4E). Interestingly, H3K4me3 signals gradually decayed in all classes despite at differential decelerating rates along pseudotime. However, H3K27me3 signals first quickly increased and plateaued in class 1. Class 2 exhibited a gradual increase in transcription, but with H3K27me3 increasing first and then slowly decreasing. Surprisingly, similar to the H3K27me3 pattern in class 2, class 3 displayed a gradual decrease in transcription along pseudotime. Likewise, we examined these dynamic patterns on constitutive genes in three quantiles ranked by the transcription level, finding no change in each molecular level along pseudotime and suggesting the specificity identified from most differentially expressed genes (Figures 4F and 4G). Together, our data indicated that H3K4me3 and H3K27me3 played context-specific roles in gene transcriptional regulation during cell fate dynamics, and that the current link between H3K4me3 and transcription may not in general support the instructive role.

### Reconstruction of single-cell histone bivalency dynamics in mESCs

Until now, simultaneously measuring more than two histone marks in single cells has been unattainable. We attempted to fill this gap through integrating separate CoTECH experiments with one on one histone mark together with transcriptome. We reasoned that single cells were subjected to CoTECH for RNA and one histone mark per experiment but multiple marks in single cells can be derived through the shared transcriptome. Towards this goal, we established an analytical framework to explore the relationship of multiple histone marks and transcriptome. By merging single cells with the closest transcriptome but with different histone marks from separate CoTECH experiments, a “pseudosingle cell” can be derived,. Histone bivalent domains have been defined as the regions co-enriched for H3K4me3 and H3K27me3, mostly around developmental genes (Bernstein et al., 2006; Harikumar and Meshorer, 2015). Using this framework, we analyzed the bivalency dynamics in mESCs (Figure 5A; STAR Methods). In general, we merged 20 single cells into one pseudosingle cell, and integrated the omics information from H3K4me3, H3K27me3 and gene expression. Second, we defined a bivalency score (Dainese et al., 2020) by calculating H3K4me3 and H3K27me3 signals in the pseudosingle cells using the equitation detailed in STAR Methods. We first tested the feasibility of this bivalency score by using bulk data. Genes marked with both H3K4me3 and H3K27me3 displayed a high bivalency score (e.g. *Drd5*) while genes marked with either one presented a low bivalency score (e.g. *Halr1* and *Hotair*). This finding was also cross-verified as genes with high bivalency scores were largely overlapped with the previously established bivalent gene list (Bernstein et al., 2006) (Figures S5A and S5B). Next, we examined the bivalency dynamics during transition from naïve to primed state. Single cells were ordered along pseudotime and 603 genes with most variable bivalency scores were clustered into three groups: decreased (cluster 1), transiently noticeable (cluster 2) and increased (cluster 3) (Figures 5B–5D). However, the expression of most genes was not yet evidently detected at this timepoint (Figure 5C). Gene Ontology term enrichment analysis revealed that cluster 1 was linked with G-protein coupled receptor signaling pathway. Cluster 2 was related to the regulation of B cell differentiation. Cluster 3 was involved in functions related to embryonic morphogenesis and organism growth (Figures 5D and 5E). This finding was also verified by specific inspection of the dynamics of the bivalency score, H3K4me3, H3K27me3, RNA around representative genes including two pluripotency genes along pseudotime (Figures 5F and S5C).

**Figure 5.**
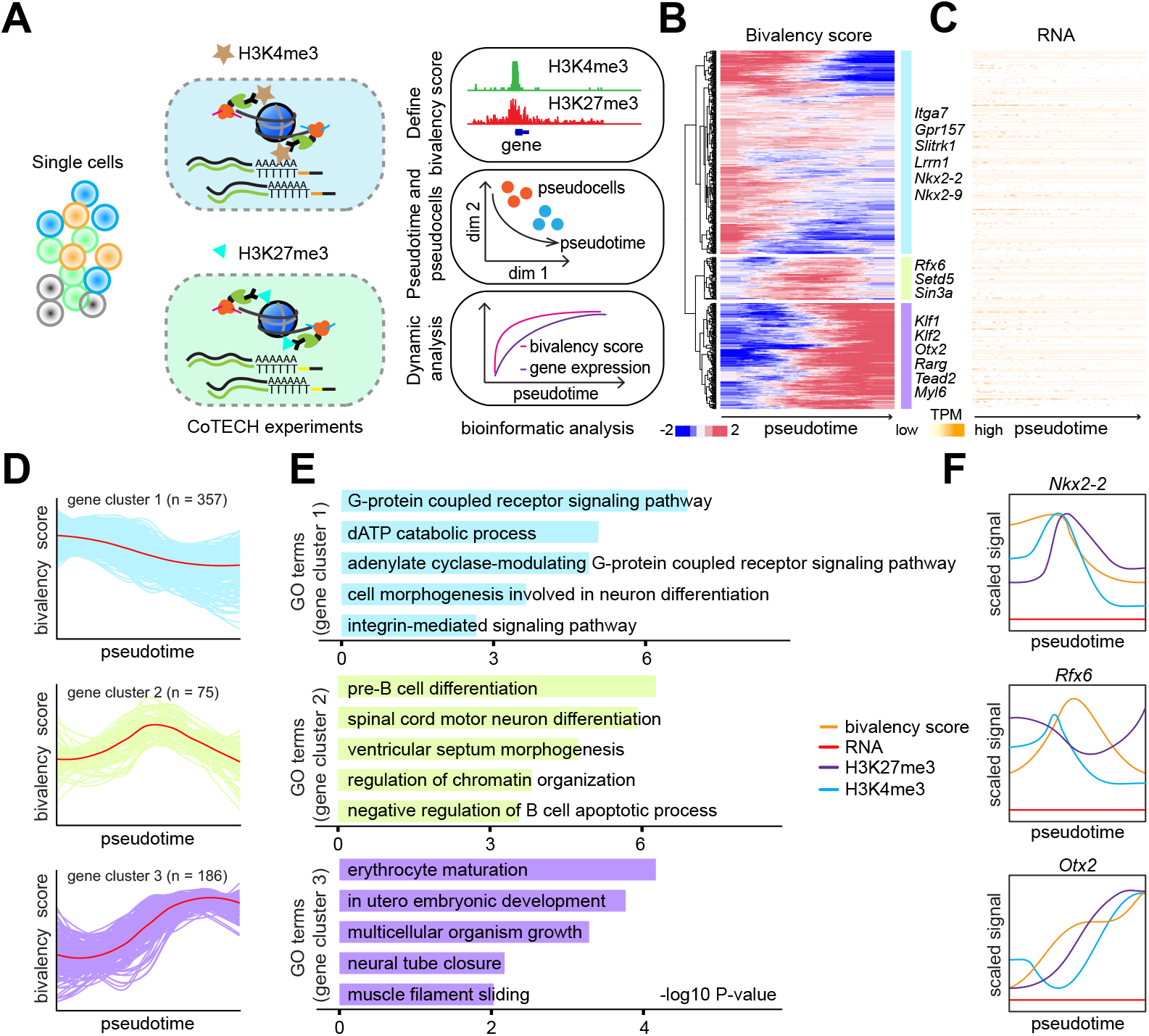
Single-cell reconstructions of dynamic trajectories of chromatin bivalency and transcription from naive to primed mESCs. (A) The framework for pseudo-single-cell reconstructions of bivalent chromatin modifications. The pseudotemporal trajectory of chromatin bivalency (H3K4me3 and H3K27me3) and gene expression in pseudo-single cells was established by using single-cell transcriptome as the anchor. (B) Heatmap showing the bivalency score of genes in three groups. Representative genes are listed alongside the heatmap. (C) Heatmap showing the expression level of genes as identified in (B) across single cells. (D) Line plots showing the dynamic patterns of chromatin bivalency in genes from three groups. Red line indicates the average score in each group. (E) GO term functional enrichment (Biological Processes) of three gene clusters as in (B). (F) Examples for the dynamic changes of H3K4me3, H3K27me3, RNA, and the bivalency score at representative genes along the pseudotime.

### CoTECH identifies differential mechanisms in cell fate decisions during endothelial-to-hematopoietic transition in two waves of hematopoietic cells

We applied CoTECH to study complex biological processes *in vivo*. Hemogenic endothelium in yolk sac (YS) contributes to Erythroid-Myeloid Progenitors (EMP) lineages as the pro-definitive wave of hematopoietic cell formation. As the definitive wave, the Aorta-Gonad-Mesonephros (AGM) is the major source of self-renewal hematopoietic stem cells (HSCs) through endothelial-to-hematopoietic transition (EHT) in the early embryo (Dzierzak and Bigas, 2018; Hou et al., 2020; Lilly et al., 2016; Palis, 2016). To understand the difference in the regulatory basis leading to programmed cell fate decisions in EHT between AGM and YS, we performed CoTECH for H3K27ac (an active enhancer mark)(Creyghton et al., 2010) and RNA in CD31^+^ endothelial cells enriched by fluorescence-activated cell sorting (FACS) in AGM and YS of E9.5 and E10.5 mouse embryos (Figure 6A). We obtained a total of 512 AGM cells and 220 YS cells passing quality control filter for subsequent analysis (STAR Methods).

**Figure 6.**
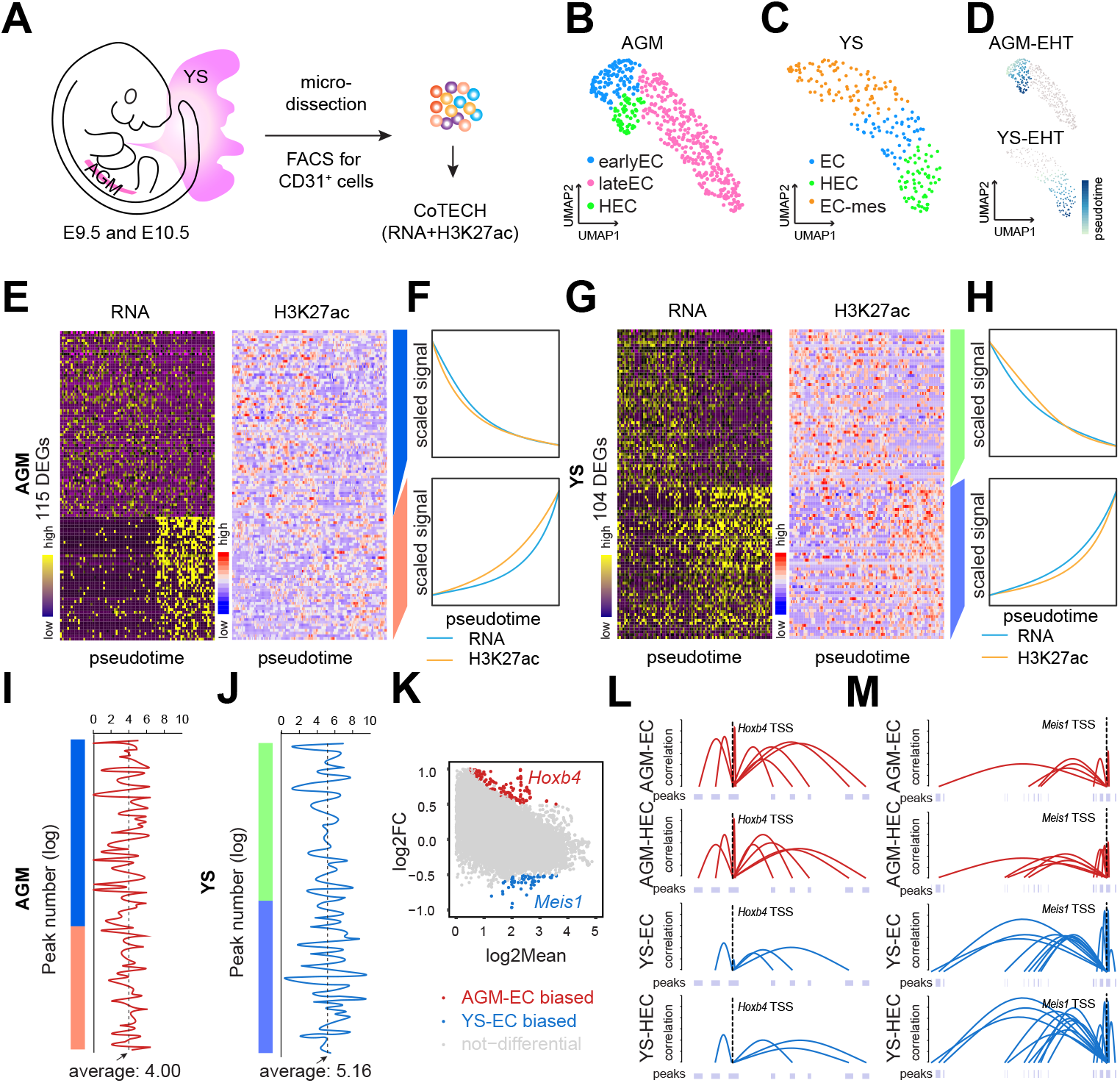
Linking enhancers to regulated genes reveals mechanisms underlying endothelial-to-hematopoietic transition in YS and AGM. (A) Experimental design using CoTECH for analyzing chromatin regulation of gene expression during endothelial-to-hematopoietic transition (EHT) from yolk sac (YS) and aorta–gonad–mesonephros (AGM) tissues. (B-C) UMAP projection based on the MOFA factors inferred using 512 AGM cells (B) and 220 YS cells (C). (D) Pseudotime anslysis of AGM-EHT and YS-EHT. (E) Heatmaps showing RNA (left) and H3K27ac (right) signals at gene-correlated peaks in 175 AGM cells. The rows represent 115 DEGs. (F) Line plots showing average RNA, H3K27ac signals of two gene clusters as in (E). (G) Heatmaps showing RNA (left) and H3K27ac (right) signasl at gene-correlated peaks in 122 YS cells. The rows represent 104 DEGs. (H) Line plots showing average RNA, H3K27ac signals of two gene clusters as in (G). (I) Distribution of peak numbers of genes shown in (E) around TSS ± 500 kb of genes. (J) Distribution of peak numbers of genes shown in (G) around TSS ± 500 kb of genes. (K) MA plot (log-ratio by mean average) comparing H3K27ac profiles between YS-EC and AGM-earlyEC. H3K27ac signals were calculated by counting the reads at the genes ±100-kb regions and normalized by read depth. (L-M) Examples for inferred enhancer-to-target gene links for Hoxb4 (L) and Meis1 (M) in YS-EC, YS-HEC, AGM-EC, and AGM-HEC.

To define cell identities or states by a higher-dimensional reconstruction of two molecular layers of single-cell data, we used multi-omics factor analysis (MOFA) (Argelaguet et al., 2018) to analyze single-cell H3K27ac and transcriptome profiles of AGM and YS cells. Both transcription and H3K27ac data of the same single cells were used as input for unsupervised dimensionality reduction in MOFA analysis. Having performed cell clustering followed by the pseudotime ordering, we identified different subpopulations annotated by specific marker genes’ expression: early EC (*Pecam1, Pf4, Mest*), late EC (*Pecam1, Cald1, Egfl7*), and HEC (*Pecam1, Itgb3, Runx1*) for AGM cells, and EC (*Pecam1, Cdh5, Ccnd2*), HEC (*Pecam1, Itgb3, Runx1*), and EC-mes (EC with mesenchymal features) (*Pecam1, Col1a1, Col1a2*) in YS cells (Figures 6B, 6C and S6A-S6F). Pseudotime analysis evidenced the transition from the endothelial to hematopoietic state during lineage specification in both AGM and YS (Figure 6D). The dynamic trend in both gene expression and H3K27ac signals at the corresponding gene-enhancer linked regions was well correlated along pseudotime, indicating the pivotal role for lineage-specific enhancers in regulating cell fate decisions during EHT (Figures 6E–6H). The EHT in YS and AGM was accompanied by the dynamic change of a set of core TFs (such as RUNX1 and SPI1) whose motif enrichment scores were calculated by chromVAR (Schep et al., 2017) in accordance with corresponding TF genes, expression (Figures S6G-S6J). We hypothesized that different regulatory mechanisms underlying transitioning in two waves of hematopoietic formation may explain the AGM-EHT path leading to definitive HSCs with potentials to reconstitute full blood lineages. Interestingly, the average number of enhancer peaks regulating corresponding target genes in YS-EHT cells was significantly higher than that in AGM-EHT cells at ± 500 kb around the differentially expressed genes (DEGs) (*P* value = 3.3e-4, Mann-Whitney test) (Figures 6I and 6J). We further identified AGM-EC biased, YS-EC biased, and common enhancer regions defined by the H3K27ac signals around the ± 100 kb of genes. *Hoxb4* is known to promote the acquisition of an HE identity and, thus, enforces early definitive hematopoietic processes (McGrath and Palis, 1997; Teichweyde et al., 2018). Consistently, we found that the H3K27ac signals around *Hoxb4* in AGM was significantly upregulated in AGM than that in YS (false discovery rate (FDR)<0.05 and log2 fold change (FC)>0.5), while the *Meis1* gene loci exhibited a higher enhancer activity in YS (Figure 6K). Next, we linked the target genes *Hoxb4* and *Meis1* to the regulatory peaks based on the covariance of H3K27ac signals and gene expression across co-assayed single cells. The inferred enhancer-*Hoxb4* and enhancer-*Meis1* pairs displayed highly selective enrichment in AGM-EHT and YS-EHT, respectively (Figures 6L and 6M). *Runx1*, an important regulator of hematopoiesis, is required for the formation of HSC and YS-derived EMPs from hemogenic endothelium during embryogenesis (Gao et al., 2018; Palis, 2016; Yzaguirre et al., 2018). Likewise, evident enhancer-*Runx1* links were identified both in AGM-EHT and YS-EHT cells, whilst more profound in YS-EHT at this stage (Figure S6K).

## DISCUSSION

In sum, we present CoTECH, a method that enables the simultaneous detection of gene expression and chromatin occupancy in thousands of single cells per experiment. CoTECH allowed assembling bivalent histone marks H3K4me3 and H3K27me3 in pseudosingle cells to delineate dynamic bivalency scores from the naïve to primed mESCs. Applying CoTECH to understanding of cell fate regulation in complex biological processes *in vivo*, we demonstrated that AGM-EHT and YS-EHT exhibit distinct underlying gene programs, which are further regulated by different enhancer-target gene links with more enhancers per gene in YS-EHT cells. This finding may have an important implication in deciphering HSC cell fate decisions and directing *in vitro* HSC reprogramming strategy for potential regenerative medicine.

CoTECH is easy-to-use for general biomedical laboratories with following advantages: First, the throughput can be scaled up to far greater numbers by simply redistributing cells to many 96- or 384-well plates in secondary indexing step; Second, no specialized devices are required to carry out CoTECH experiments, whose libraries can be readily sequenced on the standard sequencing recipe at the low cost, obviating the custom sequencing inaccessible to most laboratories. Third, CoTECH enables RNA and parallel processing of multiple histone marks that can be distinguished by different barcodes in single experiment, thus eliminating the potential batch effect;

A recent high-throughput method, scCC-seq, was developed for joint profiling of transcriptome and transcription factor binding sites (Moudgil et al., 2020). Unlike scCC-seq requiring a transgene expressing the transposase in fusion with a TF gene of interest, CoTECH is able to capture histone modifications as well as endogenous DNA binding proteins in principle, presenting more than two orders of magnitude in the DNA reads per cell than the former. Similar to scCC-seq, the method scDam&T-seq also relies on a transgenic expression of protein of interest, although it produces the comparable numbers in DNA and RNA reads per cell (Rooijers et al., 2019). However, scDam&T-seq is intrinsically at the lower throughput as based on physical isolation of single cells in 96- or 384-well plates. As with comparable quality and throughput in both DNA and RNA parts with sci-CAR in joint profiling of chromatin accessibility and gene expression, CoTECH can be easily applied to analyze complex tissue samples in patients.

We foresee that CoTECH finds wide applications in studying gene regulation and cell fate decisions. Gene expression is not the only factor that contributes to cellular heterogeneity. Epigenetic factors such as histone modifications, nucleosome occupancy, DNA methylation and chromatin conformation architecture also play active roles in transcriptional regulation and cellular heterogeneity (Efremova and Teichmann, 2020; Eisenstein, 2020; Kelsey et al., 2017; Schier, 2020; Shema et al., 2019; Zhu et al., 2020). CoTECH provides an opportunity for reconstruction of multimodal omics information in pseudosingle cells. It is imprecise to link two populations defined by different omics data. CoTECH makes it possible for integrating multilayers of molecular profiles as a higher-dimensional regulome for accurately defining cell identity (Chen et al., 2018). In this study, we exemplified in-silico assembly of bivalent histone marks in pseudosingle cell with the transcriptome as an anchor, demonstrating the single-cell resolved bivalency trajectory during the naïve to primed pluripotency transition. We believe that this can be applied to unfold the regulatory relationship between many histone modifications and chromatin binding proteins.

As with existing technologies for profiling multimodal omics in single cells, CoTECH data are still sparse. In the future, this will be improved through differently designed biochemistry. Similar to SNARE-seq, CoTECH can also be employed in the droplet-based procedure to reduce the hands-on time. Nevertheless current limitations, CoTECH provides a valuable tool for using multiomics molecular information to define cell fate determinants and potentially assemble a compendium of histone modifications and chromatin binding proteins. We further envision that CoTECH is useful to identify the rare but unannotated cell types for functional validation.

## SUPPLEMENTAL INFORMATION

Supplemental information can be found online.

## ACKNOWLEDGMENTS

We thank all members of the He lab for critical comments on this manuscript, A.H. was supported by the National Basic Research Program of China (2019YFA0801802 and 2017YFA0103402), the National Natural Science Foundation of China (31571487, 31771607 and 31327901), the Peking-Tsinghua Center for Life Sciences, and the 1000 Youth Talents Program of China.

## AUTHOR CONTRIBUTIONS

A.H. conceived and designed the study. Y.L. and Q.W. designed and performed all experiments, assisted with X.Y. H.X. performed the computational analyses, supervised by A.H. H.X., Y.L. and A.H. wrote the paper with input from all other authors. All participated in data discussion and interpretation.

