## Supplemental information for "Single-cell joint detection of chromatin occupancy and transcriptome enables higher-dimensional epigenomic reconstructions"

### METHODS

#### Mice

All mice experiments were performed according to the protocol (#IMM-HeAB-1) approved by the Institutional Animal Care and Use Committee of Peking University.

#### Cell culture

All cell lines were cultured at 37°C with 5% $CO_2$ and were maintained in high glucose DMEM (Hyclone) for HEK293T and NIH 3T3 cells or RPMI 1640 (Gibco) for K562 cells, all supplemented with 10% FBS (Sigma) and 1% Penicillin/Streptomycin (Hyclone). Wild-type V6.5 murine embryonic stem cells (mESCs) were cultured on 0.1% gelatin-coated plates in ESC DMEM culture medium containing 15% fetal bovine serum (Sigma), 1% Penicillin/Streptomycin (Hyclone), 1% Glutamax (GIBCO), 0.1 mM 2-mercaptoethanol (Sigma), 1% MEM nonessential amino acids (Cellgro), 1% nucleoside (Millipore), and 1,000 U/ml recombinant leukemia inhibitory factor (LIF) (Millipore).

#### Mouse embryonic EC preparation

Wild-type C57BL/6 mice were interbred and dissected at E9.5 and E10.5 stage. Yolk sac (YS), aorta/gonad/mesonephros (AGM) and body region of the embryo were dissected as previously described (Zeng et al., 2019). Cells were digested with 0.1% Collagenase I (Sigma) and incubated at 37°C for about 30 min. After wash once by 1% BSA/PBS, cells were stained by anti-CD31-PEcy7 (BioLegend, 102524) and incubated on ice for 30 min. Cells were washed once by 1% BSA/PBS and resuspended by 1% BSA/PBS, and PEcy7 positive endothelial cells were sorted using Beckman Coulter MoFlo XDP.

#### Barcoded PAT preparation

The procedure of PAT expression, purification, assembling was followed as previously described (Picelli et al., 2014; Wang et al., 2019). In short, pET28a-His-pA-Tn5 was transformed into BL21 (DE3) competent cell, and colonies were picked into final 500 mL LB medium, until O.D. reached 0.5-0.8. PAT expression was induced by adding 0.2 mM IPTG, and bacteria culture was incubated at 23°C, 80 rpm for 5 h. Bacteria pellets were collected and lysed by HGX buffer before sonication. Genomic DNA in the lysate was precipitated by PEI (Sigma) and clear lysate was applied to the Ni-NTA column (Qiagen). After wash, PAT was eluted from column by 100 mM imidazole. Concentration of PAT was quantified by Coomassie Blue staining after resolved in a 7.5% PAGE-gel. Both PAT and annealed barcoded adaptor were mixed at the equal molar ratio at 37.5 μM and incubated at 25°C for 60 min. The barcoded PAT can be stored at -20°C and ready to use.

**CoTECH experimental procedure**

Digested single cells were fixed by 90% methanol and stored at -80°C. Cells were collected at 1000 G, 4°C for 5 min and resuspended by Wash Buffer before incubated with 3-10 μl activated Concanavalin A (ConA) beads (Bangs Lab) at room temperature for 15 min. Samples were resuspended in 100 μl antibody buffer with addition of 0.5 μg primary antibody, and incubated at 4°C for 3 h . Barcoded PAT was added to each well, followed by incubation at 4°C for 1 h. Targeted tagmentation was carried out at 25°C for 1 h, and reverse transcription (RT) was performed using SuperScript IV (Invitrogen). After RT, all cells were combined to a 1.5 ml Eppendorf tube and stained with DAPI, and 20 cells were sorted into each well in 96-well plates containing 3 μl Elution Buffer. The plates were incubated at 55°C for 6 h and then 85°C for 15 min. Next day, samples were supplemented with 1 μl 1.8% TritonX-100 and incubated at 55°C for 5 min to quench SDS. To each well, 20 μl PCR mix was added, followed by 16 cycles of amplification. DNA was purified by 1.8X AMPure XP beads, and eluted by 10.5 μl ddH2O. 5 μl DNA elution was transferred to a new plate for the DNA-part library preparation as well as 5 μl for the RNA-part library preparation.

**DNA-part library preparation**

The 2-round PCR for the DNA-part library was performed as previously described, resulting in the standard Illumina Truseq construct (Wang et al., 2019). Briefly, 15 μl PCR mix (0.2 μl KAPA HiFi HotStart DNA Polymerase, 4 μl 5 x KAPA High-GC Buffer, 1 μl 10 mM dNTP Mix, 0.4 μl 25 mM $MgCl_2$, 8.9 μl $H_2O$, and 0.5 μl 50 μM connector A/B primer mix) was added to each well. PCR was performed as following: 1 cycle of 95°C 3 min; 8 cycles of: 98°C 20 s, 65°C for 30 s, 72°C for 1 min; 1 cycle of 72°C for 5 min, hold at 4°C. Excessive primers were digested by adding 0.25 μl ExoI (NEB) and incubated at 37°C for 60 min followed by 72°C for 20 min. 8 μl of second round PCR mix (0.2 μl KAPA HiFi HotStart DNA Polymerase, 2 μl 5 x KAPA High-GC Buffer, 0.2 μl 25 mM $MgCl_2$, and 5.6 μl $ddH_2O$) was added to each well containing 1 μl 10 mM Truseq index i5 and 1 μl Truseq index i7, and subjected to amplification: 1 cycle of 95°C 3min; 5 cycles of 98°C for 20 s, 65°C for 30 s, 72°C for 1 min; 1 cycle of 72°C for 5 min; hold at 4°C. Samples were pooled and purified by the TIANquick Mini Purification Kit (TIANGEN) and 0.8 × AMpure XP beads (Beckmann). Size selection was also performed by the gel extraction for the fragments at the size of 400-1,000 bp on 1.5% agarose gel. DNA was finally purified by 0.8 × AMpure XP beads (Beckmann) and ready for sequencing.

**RNA-part library preparation**

For each well, 2.5 μl tagmentation mix (0.75 μl 10X TAPS-MgCl2, 0.75 μl DMF, 1 μl Tn5-MEA) was added and reaction was performed at 55°C for 5 min. After tagmentation, 1 μl 0.2% SDS, 0.4 μl BSA (NEB) was added and incubated at 55°C for 5 min. To quench SDS, 1 μl 1.8% TX-100 was added and incubated at 55°C for 5 min. 28.1 μl of PCR mix (0.4 μl KAPA HiFi HotStart DNA Polymerase, 8 μl 5 x KAPA High-GC Buffer, 1 μl 10 mM dNTP Mix, 0.8 μl 25 mM $MgCl_2$, and 17.9 μl $H_2O$) was added to each well containing 1 μl 10 mM Nextera index i5 primer and 1 μl Truseq index i7 primer. The PCR enrichment was performed in a thermal cycler: 1 cycle of 72°C for 5 min; 1 cycle of 95°C for 3 min; 11 cycles of 98°C for 20 s, 65°C for 30 s, 72°C for 1 min; 1 cycle of 72°C for 5 min, hold at 4°C. Product was pooled and purified by TIANquick Mini Purification Kit (TIANGEN) and 0.8 × AMPure XP beads (Beckmann). Size selection was also performed by the gel extraction for the fragments at the size of 400-1,000 bp on 1.5% agarose gel. DNA was finally purified by 0.8 × AMpure XP beads (Beckmann) and ready for sequencing.

**Data processing**

**CoTECH data processing-scRNA-seq part**

First, the CoTECH RNA-part data were demultiplexed using in-house scripts with tolerating one mismatched base in each 8-base barcode. Raw sequencing reads with adapters and low-quality based were trimmed by cutadapt (version 1.11). The read 1 sequence were mapped to the reference genome (mm10 and hg19 for mouse and human sample, respectively) using Hisat2 (version 2.0.4) (Kim et al., 2015) with single-end mode. Uniquely mapped reads were counted using the package HTSeq (Anders et al., 2015). The gene expression level was quantified by the UMI number of each gene normalized by the total UMI numbers.

**CoTECH data processing-DNA part**

The CoTECH DNA-part data were demultiplexed by custom scripts. Single cell libraries were processed to generate unique and non-duplicated libraries as previously described (Wang et al., 2019). Briefly, DNA sequencing data was evaluated by FastQC (version 0.11.5). Adapters and low-quality bases at the end of sequence reads were removed using cutadapt (version 1.11) with the following parameters: -q 20 -O 10 --trim-n -m 30

--max-n 0.1. Paired-end ChIP reads were then mapped to reference genome (mm10 and hg19 for mouse and human samples, respectively) using Bowtie2 (version 2.2.9) (Langdon, 2015). Uniquely mapped reads were obtained and sorted by Samtools (version 1.9). Duplicated reads from the PCR amplification step were identified and removed by Picard (version 2.2.4) (http://broadinstitute.github.io/picard) with default parameters. We used MACS2 (version 2.1.1) (Zhang et al., 2008) to call peaks based on the uniquely mapped, non-duplicated reads.

**Calculating collision rate**

To evaluate the efficiency of combinatorial cellular indexing for single-cell labelling, we calculated the collision rate, i.e., the ratio of two cells or more coincidentally labelled by the same combination of barcodes. We performed H3K27ac CoTECH experiments using the 1:1 mixture of human (HEK293T) and mouse (NIH 3T3) single cells. We plotted the scatterplot using proportion of human reads in each barcode combination in Figure 2. Barcodes with fewer than 80% of aligned reads mapped to one species were classified as collisions (Murschhauser et al., 2019).

**Visualization and correlation analysis of CoTECH data**

We used bamCoverage in deepTools (version 2.2.3) (Ramírez et al., 2014) to calculate coverage in bam files and generated track files (bigwig format) for CoTECH DNA and RNA data. For visualization of the DNA and RNA reads at specific loci, the bigwig files were uploaded into IGV (version 2.3.59) (Thorvaldsdóttir et al., 2013) together with reference tracks.

For correlation analysis between different experiments in Figure 2 and Figure S4, we quantified DNA signals at peak regions by RPKM (reads per kilobase per million mapped reads). RNA was normalized to TPM (transcripts per million). For single cells with both H3K4me3 and transcriptome data in Figure 4, the correlations between H3K4me3 signals at TSS±5 kb or gene body±100 kb and gene expression were assessed. Pearson correlation coefficients were calculated between each comparison.

**Comparison with published single-cell multi-omics approaches**

We benchmarked CoTECH data by the non-duplicated reads (DNA part), UMI number (RNA part) and gene number (RNA part) with published single-cell multi-omics technologies that otherwise simultaneously profile open chromatin and transcriptome in the same cell (sci-CAR, Paired-seq, and SNARE-seq (Cao et al., 2018; Chen et al., 2019; Zhu et al., 2019)). The published datasets were downloaded from GSE117089, GSE130399, GSE126074.

**Clustering of CoTECH data in mESCs**

We first identified cells that both DNA and RNA profiles were obtained based on the matching barcode combinations in the four mESC H3K4me3/H3K27me3-RNA CoTECH experiments (Cells incubated with either H3K4me3 or H3K27me3 antibody were distributed into different wells in 96-well plates and distinguished by specific indexing). We filtered out cells with fewer than 500 non-duplicated reads and 3,000 UMIs and removed top 2% cells with an atypically large quantity of UMIs and non-duplicated reads, as these cells may represent doublets (Ai et al., 2019; Fan et al., 2018), resulting a total of 6,993 cells for subsequent analyses.

For RNA part of mESC CoTECH data, we used Seurat (version 3.0.2) (Satija et al., 2015) to identify cell types. Genes detected in at least 10 cells were kept for further analysis. We performed PCA with top 2,000 highly variable genes (HVGs) selected using “FindVariableGenes” function. Top 20 significant PCs were used to perform UMAP analysis and cluster identification using the “RunUMAP” and “FindClusters” functions, respectively. We identified two clusters that were further annotated by known marker genes, and 129 differentially expressed genes (DEGs) between the cell types were identified with the function “FindAllMarkers” in the R package Seurat. The mESCs were ordered in pseudotime by Monocle 2 package (version 2.6.3) (Qiu et al., 2017) as described in the tutorials (http://cole-trapnell-lab.github.io/monocle-release/). HVGs identified by Seurat were used as input for temporal ordering of these cells. The remaining parameters in Monocle 2 were in default.

To reduce the dimension and identify subpopulations by DNA data, we used cisTopic (Bravo Gonzalez-Blas et al., 2019) to perform single-cell analysis on H3K4me3 (3,907 cells) and H3K27me3 (3,086 cells). We called peaks for aggregated DNA profiles in each cluster defined by RNA profiles, and cluster-specific peaks were used as input to run cisTopic with default parameters. We visualized the cell states using a dimensionality reduction method Diffusion map in R. A density-based method (densityClust package in R) was used to perform cell clustering (Preissl et al., 2018).

**Chromatin bivalency analysis**

We provided an analyze framework to analyze the dynamic change of multiple histone marks at single-cell level. Here, we focused on the bivalent features, i.e. genomic loci with both H3K4me3 and H3K27me3 modification, in mESC as an example. To reduce the sparsity of CoTECH DNA data, we merged 20 cells into one “pseudocell” based on the similarity of RNA profiles, resulting in 350 pseudocells for mESC CoTECH data. H3K4me3 and H3K27me3 ChIP signal of each pseudocell were calculated using the normalized read intensity of H3K4me3 and H3K27me3 ChIP at the ±100 kb regions of genes. To accurately study the dynamics of bivalency along pseudotime, we defined the bivalency score for gene *i* in the pseudocells as:

$$bivalency\ score_i = \frac{K4i + K27i}{abs(K4i - K27i) + 1}$$

where *K4i* and *K27i* are the normalized read intensity in gene *i* for H3K4me3 and H3K27me3. Total 603 genes with most variable bivalency score were clustered to 3 groups, and the bivalency score of genes were visualized by heatmap using the R package ComplexHeatmap (version 2.0.0). Gene ontology analysis in Figure 5 was performed using DAVID (https://david.ncifcrf.gov/home.jsp) (Huang et al., 2009a, b).

**Data analysis of the hematopoietic lineage during endothelial-to-hematopoietic transition**

To study epigenomic regulation of the transcription program of endothelial-to-hematopoietic transition occurring in extraembryonic yolk sac and AGM regions, we first identified subpopulations of 512 AGM and 220 YS $CD31^+$ cells by MOFA (Argelaguet et al., 2018). MOFA was used to unsupervised dimensionality reduction by simultaneously inputting single-cell DNA and RNA-seq data. The subpopulations of YS and AGM cells were identified and annotated by marker genes. Single cells were ordered based on the UMAP coordinate and local distance in the clusters as previously described (Granja et al., 2019). To infer the regulatory relationship of enhancers to their target genes, we calculated the correlation between gene expression and the normalized H3K27ac signals of candidate peaks around the TSS of each gene from cisTopic.
